# synder: inferring genomic orthologs from synteny maps

**DOI:** 10.1101/554501

**Authors:** Zebulun Arendsee, Andrew Wilkey, Urminder Singh, Jing Li, Manhoi Hur, Eve Syrkin Wurtele

**Affiliations:** Dept. of Genetics Development and Cell Biology, Iowa State University, Ames IA, 50010, USA; Center for Metabolic Biology, Iowa State University, Ames, IA 50011, USA; Dept. of Agronomy, Iowa State University, Ames, IA 50011, USA

## Abstract

Ortholog inference is a key step in understanding the evolution and function of a gene or other genomic feature. Yet often no similar sequence can be identified, or the true ortholog is hidden among false positives. A solution is to consider the sequence’s genomic context. We present the generic program, `synder`, for tracing features of interest between genomes based on a synteny map. This approach narrows genomic search-space independently of the sequence of the feature of interest. We illustrate the utility of `synder` by finding orthologs for the *Arabidopsis thaliana* 13-member gene family of Nuclear Factor YC transcription factor across the Brassicaceae clade.

## 1 Introduction

A powerful first step in understanding the evolution and function of a genomic feature is resolving its genomic context, that is, comparing the feature to orthologous features in other species. Comparing multiple orthologous features across species allows evolutionary patterns to be uncovered. These patterns may include evidence of purifying selection, which implies the feature is important to the survival of the species; positive selection, implying the feature is rapidly evolving along one lineage; and functional dependencies between sites (for example, amino acids in an enzyme reaction site) [1]. These evolutionary trends have direct application in fields such as rational protein design [2]. Distinguishing between orthologs (homologous features arising through speciation) and paralogs (homologous features arising through gene duplication) is foundational to understanding the history of a feature. Genomic context is also critical for discerning the origins of the often large numbers of species-specific “orphan” genes that are found in most genome projects [3–6].

Identifying orthologs is not easy. A simple sequence similarity search of a query feature (e.g., a gene, transposon, miRNA, or any sequence interval) against a genome or proteome of a target species may obtain thousands of hits in a swooping continuum; these could include: the true ortholog, related family members (paralogs), and non-specific hits. Therefore, methods for winnowing the search results have been developed to identify the true orthologs. A straightforward approach to identify orthologs of protein-coding genes is reciprocal best hits [7]. In this technique, a protein encoded by a gene from the focal species is searched (e.g. with BLAST) against the target proteome. The highest scoring gene is then searched back against the proteome of the focal species. If the top scoring hit of the second search is the original query gene, then the two genes are accepted as orthologs. There are also methods that build on reciprocal best hits, such as the reciprocal smallest distance method that considers evolutionary distance in addition to similarity score [8].

Little or no significant sequence similarity is expected across species for some classes of features. A lack of significant similarity may stem from sequences being very short (e.g., a single promoter element or an miRNA) or it could result from very rapid mutation rates in the feature (e.g., intragenic intervals that are under little or no purifying selection). Orphan genes, which by definition have no protein homolog in related species, are an example of a feature for which sequence comparisons alone cannot delineate the region in the target genome from which the orphan gene arose [4]. These genes are often both short *and* rapidly evolving, making it very difficult to find orthologous genomic regions (possibly non-coding) even in closely-related target species. Without an ability to identify orthologous genomic intervals, the pathway of evolution of an orphan cannot be determined; for example, orphans of de *novo* origin cannot be distinguished from those orphans that stem from rapid mutation [3].

Purely sequence-based methods are also problematic if the true ortholog of a query gene is duplicated in the target species. In this case, the target species contains two genes that are true orthologs of the query gene. The co-evolution of duplicate genes relative to their singleton ortholog, is of interest in theoretical evolution [9]. One of the copies may rapidly evolve to gain a new function or it may become a pseudogene [10]. In either case, the reciprocal best hits method would find only the conserved copy.

A different approach to ortholog identification, one which does not depend on the genome-wide sequence similarity of the query features themselves and that does handles duplication events, is to consider the genomic context of the query, i.e., synteny [11]. Genomic synteny is the conservation of the order of genomic features between two genomes [12].

The most obvious approach to a context-based search is to include the flanking regions of a query feature of interest when searching for an ortholog in the target genome. This approach is used by MicroSyn [13] for finding orthologs of features, such as miRNAs, that are too short and numerous to be easily searched by their sequence alone. While this approach works well for an individual query feature, extending it to a high-throughput analysis is problematic, since no single cutoff for flank length will work well for all cases. For instance, a sequence residing within a highly repetitive centromere might require flanks of megabases.

An alternative to looking at the flanking sequence of each query feature individually is to reference a genome-wide synteny map. Rather than searching for the feature directly, orthologs of flanking syntenic regions (blocks) can be identified, and a potential ortholog of the query feature can be identified in the target genome by searching within the syntenic region. This strategy has been applied to study the genomic origin of orphan genes [14] (and the method refined in [15]) where a map of one-to-one orthologous genes was used to infer the orthologous genomic intervals where the non-genic sequence corresponding to the orphan genes is expected to reside. The one-to-one map made the computational problem very easy, and could effectively identify a sub-set of the genes of de *novo* origin, but the map was very coarse, especially in regions of low gene density, so no information is obtained for other orphan genes.

There are many programs designed to build synteny maps. Some programs build sparse synteny maps from given sets of orthologous genes (e.g., OrthoClusterDB [16]). Others perform full genome alignments. Of these, some focus on large scale (megabase range) syntenic blocks that are conserved across great evolutionary distances (e.g., `DiagHunter` [17]), while others focus on micro-synteny, producing maps of many small syntenic blocks that capture local inversions, duplications and deletions (e.g., `BLASTZ` [18], `MUMmer4` [19], and `Satsuma` [20]). These micro-synteny programs are of greatest interest in this paper.

The diverse synteny mapping tools, though highly variable in granularity and accuracy [11,21], provide powerful approaches to enable the study of comparative genome evolution [12,22–24] and to glean novel information about the origin of *de novo* orphan genes [14,15,25–27]. However, the use of these maps as a tool for orthology has been generally limited to either manual inspection, or to considering only those query features that overlap syntenic blocks.

`synder` is designed to infer orthologous regions in the target genome, even when the orthologs are *between* syntenic blocks, and to assess the quality of the inferences. To do this, it traces query features from a focal genome to a target genome using a whole-genome map. `synder` is a high-performance program with a core written in C++ and an R wrapper for integration into R workflows. It will work with any synteny map, but was designed for fine-grain micro-synteny maps that capture local inversions and transpositions. It assembles collinear sets of syntenic blocks from the map and uses them to infer tight search intervals for each query on the target genome, naturally handling duplication events and inversions. `synder` also provides detailed information about the quality of the search result. The only input required is a whole-genome synteny map and a set of features of interest in the focal genome. Thereby, `synder` automates the use of syntenic information to study orthologs across any pair of species with sufficiently conserved synteny.

## 2 Algorithm

The primary function of `synder` is to map a user-designated set of query features in the focal genome to a set of search intervals in a target genome (see **Table 1** for terminology and **Figure 1** for overview). To do this, `synder` contextualizes the query features based on a user-provided synteny map for the focal and target genomes. Query features of the focal genome are mapped to an associated synteny-based search interval on the target genome; this search interval delineates the region of the target genome where the query feature is predicted to be located.

**Figure 1:**
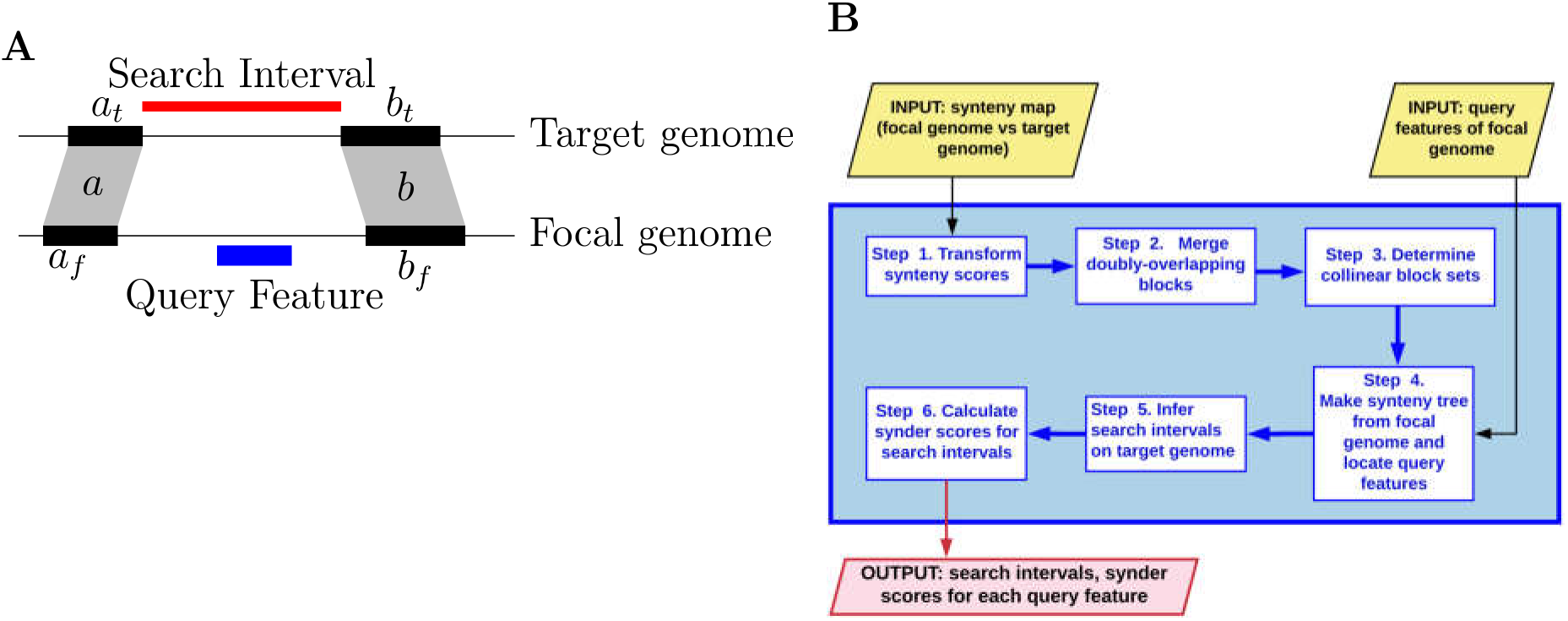
The `synder` algorithm identifies search intervals for query features based on synteny. (**A**) Diagram of a very simple syntenic relationship across focal and target genomes. *a_f_, a_t_, b_f_*, and *b_t_* are four syntenic intervals that comprise block a (*a_f_, a_t_*) and b (*b_f_, b_t_*). Blocks *a* and *b* are collinear and provide landmarks for associating the **query feature** in the focal genome with its **search interval** in the target genome. (**B**) Flow chart of the steps in the `synder` algorithm. `synder`: 1) transforms the synteny scores for each of the blocks in an input synteny map, such that scores are additive; 2) merges doubly-overlapping blocks; 3) assigns each block in the synteny map to exactly one collinear set of blocks; 4) finds the overlapping or nearest flanking syntenic intervals for each query feature in the focal genome (e.g., *a_f_* and *b_f_* in **A**); 5) for each query feature (i.e. the interval corresponding to a feature of interest on the focal genome) finds all collinear block sets that contain at least one of the blocks that flank or overlap the query feature, and then relative to each of these collinear block sets, maps the query interval to a search interval in the target genome; and 6) calculates search interval scores for each query feature relative to the search interval of each collinear set. The final output provides the query features with their corresponding hits in the target genome and a composite score for each hit.

**Table 1:** Terminology

|  |  |
| --- | --- |
| focal genome: | The genome that contains the query features. |
| target genome: | The genome in which search intervals are found for each feature of interest. |
| query feature: | The sequence interval delineating any feature of interest in the focal genome. This could be a protein-coding gene, an miRNA, an intron, a transposon, a nucleotide repeat, an lncRNA or any other genomic feature |
| blocks: | Focal and target genome intervals that are inferred to be orthologs by an outside synteny program. |
| synteny map: | A set of blocks for a pair of genomes. |
| syntenic interval: | A single interval on one side of a synteny map. |
| adjacent intervals: | Two syntenic intervals on the same scaffold with no syntenic interval located entirely between them. |
| collinear blocks: | Two blocks where both the focal and target syntenic intervals are adjacent and in the same orientation. |
| collinear block set: | An ordered set of blocks where block $i$ is collinear to block $i + 1$. |
| query context: | All blocks that overlap or are adjacent to the query interval. |
| search interval: | An expected location of an ortholog of a query feature in the target genome. |
| search space: | The union of search intervals for a given query interval. |
| synteny score: | A score for a syntenic block produced by the outside synteny program. |
| synder score: | A score for the relative reliability of a search interval (see **Figure 5**). |

**Algorithm 1** is an overview of the `synder’s` search algorithm. Each of the functions in this algorithm is defined in detail in the subsequent sections.

**Algorithm 1:**
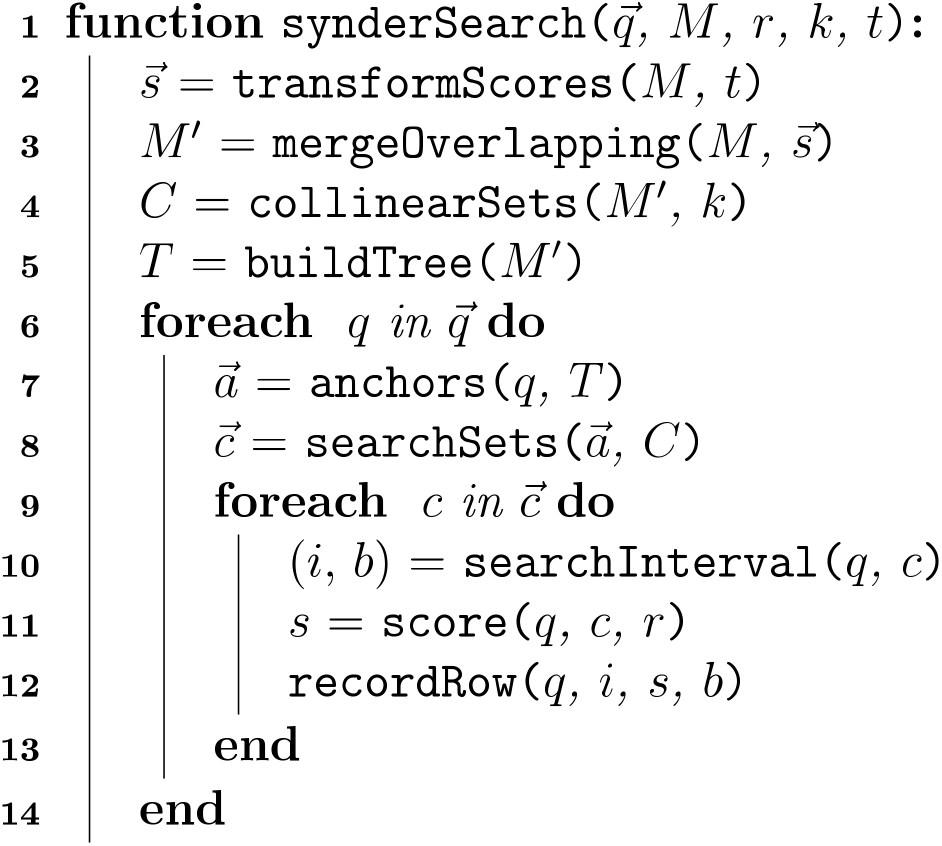
A high-level overview of the core `synder` search algorithm. 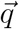, list of query features; *M*, synteny map; *r*, search interval score decay rate (see **Figure 5**); *k*, number of interrupting blocks that is tolerated; *t*, type of synteny score; 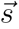, vector of transformed, additive scores used in assigning final scores to each search interval. `synder` transforms scores and merges overlapping blocks to yield a processed, reduced synteny map, *M*′. Sequential syntenic blocks, *C*, are determined from *M*′. *T*, the interval tree data structure, is then used to find the syntenic context (i.e., the anchors, 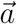) on the focal genome for each query feature, *q*. Next, query features are mapped to one or more collinear set of blocks, 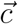. For each block, the associated search interval, *i*, is identified and the type of boundary, *b*, is determined. Each search interval is given a `synder` score, *s*. Finally, each search interval is recorded in the output table as a single row including the query feature (*q*), search interval (*i*), `synder` score (*s*), and search interval type (*b*).

### 2.1 Input Synteny Map

The primary raw input to `synder` is a synteny map that is provided as a table where each row describes one block. Each block consists of: an interval in the focal genome; an inferred syntenic interval in the target genome; a synteny score representing some metric of the confidence that the pair of intervals is orthologous; and, the relative orientation of the intervals. The focal and target intervals are each described by a chromosome/scaffold name and a start and stop position. The synteny score for each block is some measure of quality/certainty (e.g., percent identity or p-value) that is specific to the tool that used to generate the map. The orientation of the block is the strand in the target genome relative to the query, with ‘+’ indicating the same strand and ‘-’ indicating the inversion.

### 2.2 Step 1. Transform synteny scores

The synteny scores for the blocks in a synteny map may be expressed in a variety of ways by the various synteny programs. Strong similarity may be represented by low numbers (e.g., if scores are e-values) or high numbers (e.g., if scores are bitscores). Scores may be additive (e.g., bitscores) or averaged (e.g., percent similarity). The user must specify the type of the input synteny scores. Internally, the `synder` algorithm transforms these scores so that they are additive. More specifically, `synder` assumes *S*(*a* + *b*) = *S*(*a*) + *S*(*b*), that is, if the blocks *a* and b are concatenated, then the synteny score should be equal to the sums of the scores for blocks *a* and *b*. `synder` transforms the synteny map scores to an additive score using one of the transforms below:

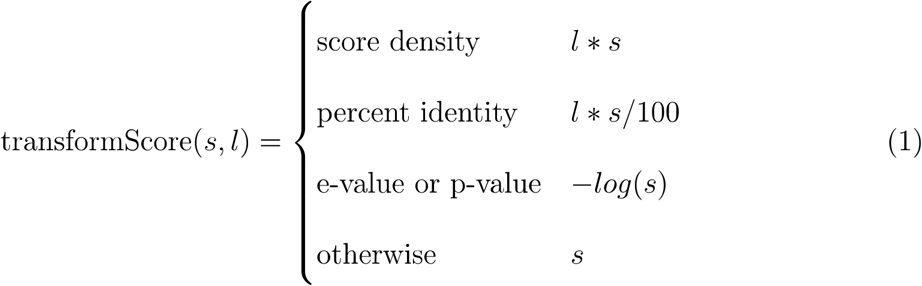

Where *s* is the input synteny score and *l* is the interval length. `synder` transforms the scores when it loads a synteny file, updates them as needed in Step 2, and ultimately uses them in Step 6 to generate scores for the final search intervals.

### 2.3 Step 2. Merge doubly-overlapping blocks

In a “perfect” synteny map, blocks would not overlap on both the focal and target sides. In practice, however, synteny algorithms occasionally produce overlapping blocks. These cases would produce multiple collinear block sets that have the same orientation and cover the same region. To avoid this, `synder` merges any blocks that overlap on both the focal and target sides. The interval of the merged blocks is the union of the overlapping block intervals. The `synder` score of the merged blocks is calculated by summing the non-overlapping interval scores with the maximum of the overlapping intervals:

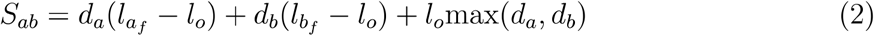

Where *d_a_* and *d_b_* are the score densities of blocks *a* and b (density is the synteny score for a block divided by the length of the syntenic interval on the focal genome); and where *l_a_f__, l_b_f__* and *l_o_* are the lengths of *a, b*, and their overlap, respectively.

A potential downside of this approach is that, when more than two intervals are doubly-overlapping, the order in which the scores are merged matters, with blocks merged later having a stronger influence. A second issue is that the merged score is calculated based on the intervals on just one side of the synteny map. The length of each interval, and the length of the overlap between the intervals, may vary between the two sides of the synteny map. For now, we do not address either of these issues, since doing so would complicate the algorithm and probably have little effect on any biological dataset (since doubly-overlapping intervals are uncommon).

The output of Step 2 is a processed synteny map without doubly-overlapping blocks.

### 2.4 Step 3. Determine collinear block sets

`synder` assigns each block in a synteny map to exactly one set of collinear syntenic blocks (**Figure 2**). Each collinear block set consists of adjacent blocks that are ordered on the query and target sides. `synder` considers two syntenic intervals “adjacent” if they are on the same scaffold and no syntenic interval is contained entirely between them. Adjacency on the target-side further requires that the intervals have the same orientation (+/−) relative to the focal genome. Two blocks are collinear if the syntenic intervals on the focal-genome and target-genome are adjacent. The collinear block sets may be inverted and/or may overlap other collinear block sets on either the focal or target side (e.g., for duplicated sequences). The individual blocks that make up the collinear set are used in Steps 4 and 5 to delimit the search intervals on the target genome relative to each query feature of the focal genome.

**Figure 2:**
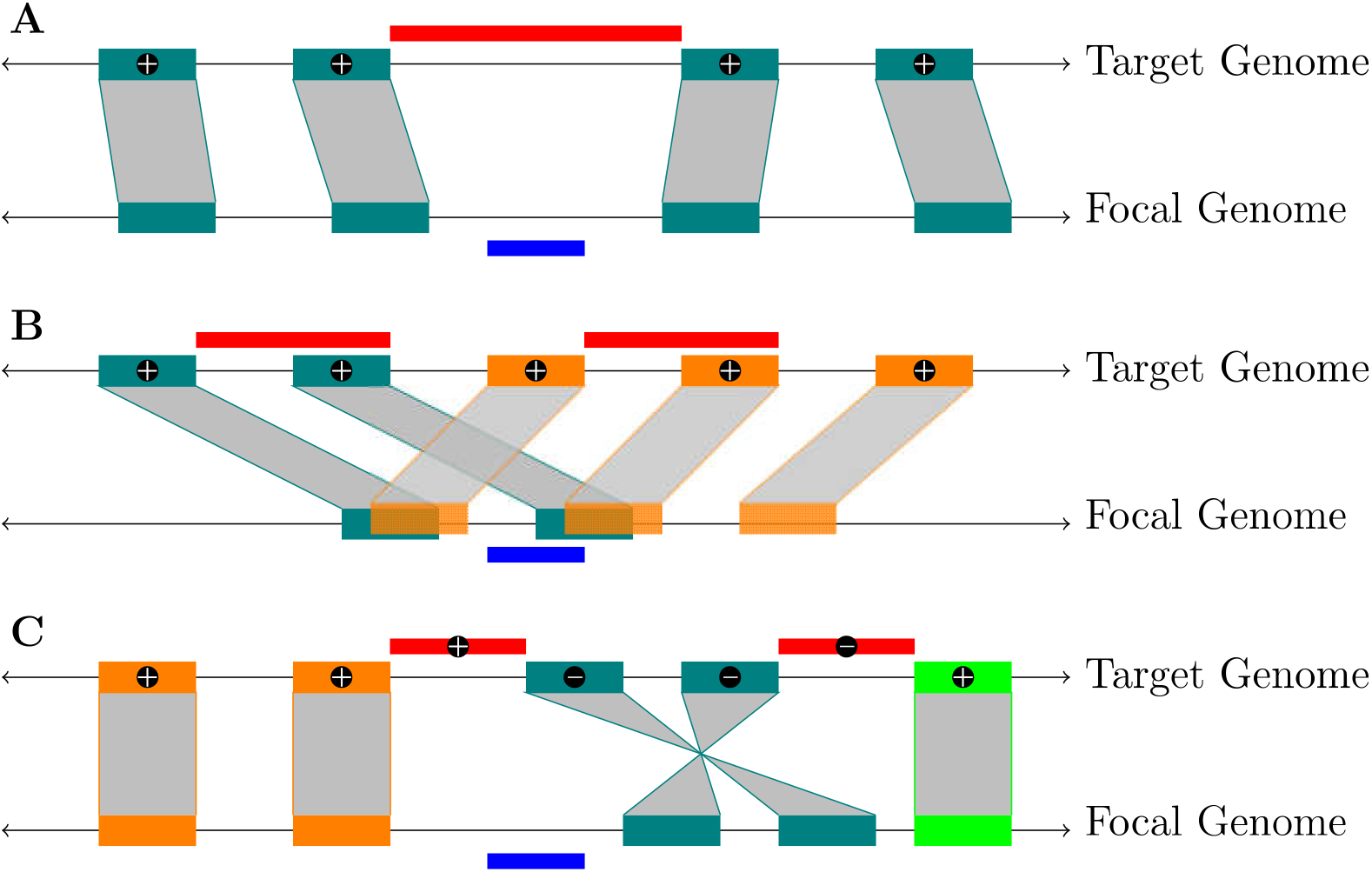
Collinear block set construction with focal-genome “anchors” to infer search intervals on the target genome. The **blue bars** below each focal genome are the query features. Genome-wide collinear block sets (colored orange, teal or green) are identified in Step 3, and are used to identify the focal-side anchors for each query feature in Step 4. The **red bars** above the target genome are the search intervals inferred by `synder` in Step 5. (**A**) a simple case where the query feature does not overlap a syntenic interval and is bound between syntenic intervals in a collinear set of blocks. (**B**) a tandem duplication where `synder` resolves the blocks into two collinear block sets (teal and orange) and infers search intervals for each. (**C**) a query feature that is **unbound** on each side (see **Figure 4**) resulting in one search interval relative to the orange collinear block set (red bar on left) and one search interval relative to the inverted teal block collinear set (red bar on right). (+/−) signs in the target search intervals represent their strand orientation relative to the query (‘−’ is an inversion).

This approach can be overly strict, resulting in many small collinear block sets. Tracing blocks across whole genome duplication, and subsequent genome alterations [28], is particularly challenging, since intervals in the homologous chromosomes could randomly diverge, resulting in a synteny map that alternates between mapping to one chromosome and the homologous chromosome. This is especially problematic in plants, where whole genome duplications are common [29]. To reduce this potential complication, `synder` provides the user an option to relax the adjacency restriction by allowing k syntenic intervals that map to alternative target scaffolds to interrupt a pair of query-side intervals in a collinear set.

The output of Step 3 is the set of blocks that are non-overlapping and adjacent. Each block is assigned to exactly one collinear block set.

### 2.5 Step 4: Find the contextual anchors on the focal genome for each query feature

The next step is to find the blocks that contain, overlap, or are adjacent to each query feature on the focal genome. These blocks will provide “anchors” that will be used in Step 5 to map the query feature to one or more collinear sets of blocks in the target genome and hence to identify the search interval(s) in the target species.

The user provides the query features as a Gene Feature Format (GFF) file that describes the genomic intervals on the focal genome corresponding to the query features of interest. A modification is to provide the GFF file along with BLAST or other whole-genome similarity scores; this modification was used as input for the case study on the NF-YC gene family (see RESULTS section).

`synder` uses a modified interval tree algorithm to locate the syntenic intervals on the focal genome that “anchor” the query feature. Building the interval tree is an *O*(*n* log(*n*)) operation (see **Algorithm 2**) and searching for a given interval is *O*(log(*n*) + *m*), where *m* is the number of overlapping intervals returned and *n* is the size of the synteny map (see **Algorithm 3**). We modified an algorithm that returns only directly overlapping intervals [30], to enable `synder` to find the flanking intervals (upstream and downstream intervals) when no overlapping intervals are found (see **Figure 3**).

**Figure 3:**
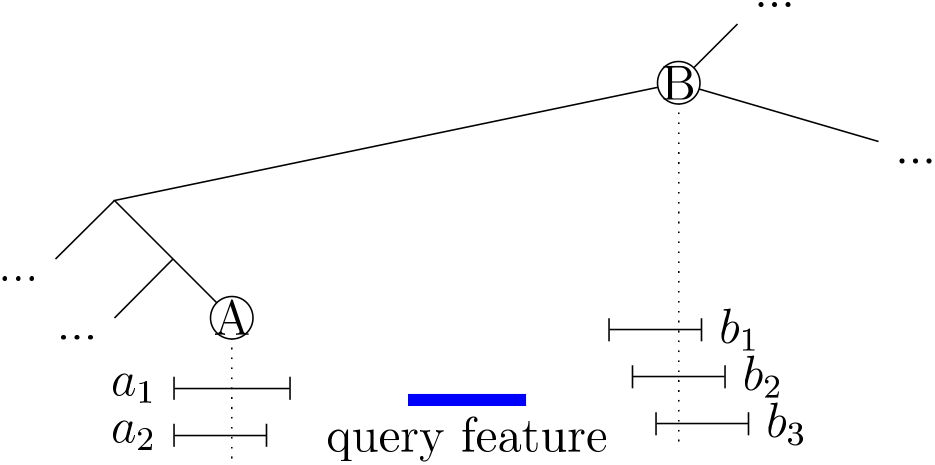
Identification of the syntenic intervals that anchor a query feature on the interval tree from the focal genome. *A* and *B* are nodes in the interval tree. *A* stores the overlapping syntenic intervals *a*_1_ and *a*_2_. *B* stores the overlapping intervals *b*_1_, *b*_2_ and *b*_3_. The query feature falls between the syntenic intervals stored in nodes *A* and *B*. The interval tree algorithm first makes the tree, and then finds the nearest node to the query. If the closest node found is *A*, and the nearest syntenic interval (*a*_1_) detected is on the left side of the query feature, then the algorithm will trace the tree until it finds the first node to the right of the query feature (i.e., node *B*). Conversely, if the closest node found is to the right of the query feature (node *B*), then the tree is traced one branch to the left, and then as many branches to the right as possible (i.e., until node *A* is found). In either case, all overlapping nodes in *A* and *B* are returned as the anchors for this query sequence.

**Algorithm 2:**
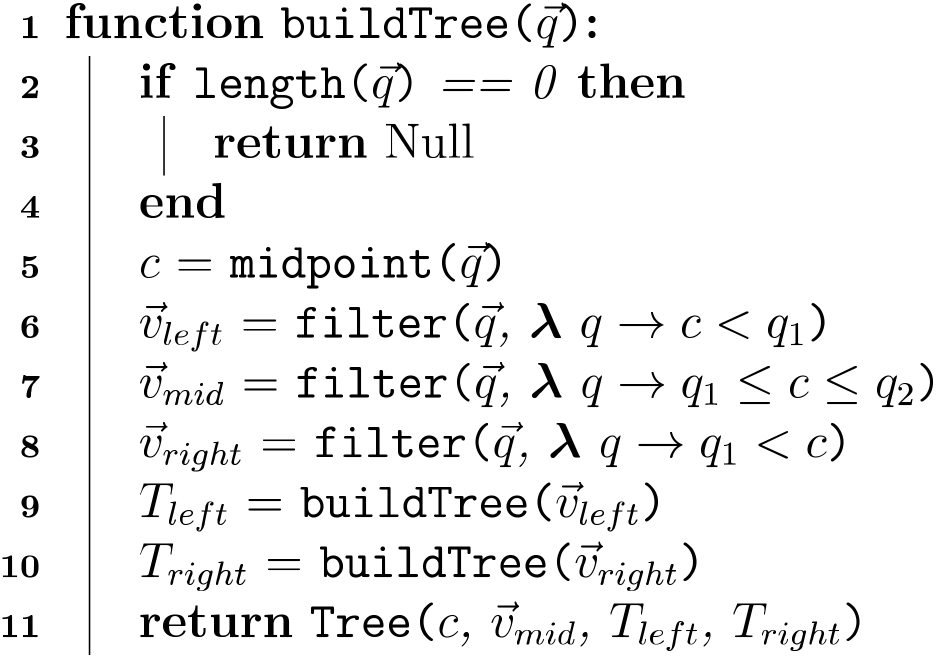
Build a synteny interval tree. buildTree takes a vector of intervals, 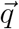, on a given scaffold/chromosome of the focal genome and returns an interval tree data structure. The midpoint c is an integer equal to the middle position in the interval in the middle of the vector of intervals (by index). If the input vector is sorted, then the midpoint will tend to be near the center of the scaffold. 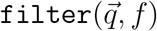 selects the subset of intervals in 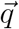 for which the condition *f*(*q*) is true. The filters in lines 6-8 partition each element in 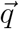 into one of three sets: intervals on the left of the midpoint *c*, intervals overlapping the midpoint *c*, and intervals on the right of the midpoint *c*. New trees are created recursively for the left (less than) and right (greater than) sets of intervals. `buildTree` returns a new syntenic interval Tree object, (*T*), that stores the midpoint c, all overlapping syntenic intervals 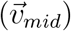, and the left and right child trees. The Tree will be used in **Algorithm 3** to identify the anchors for each query feature.

**Algorithm 3:**
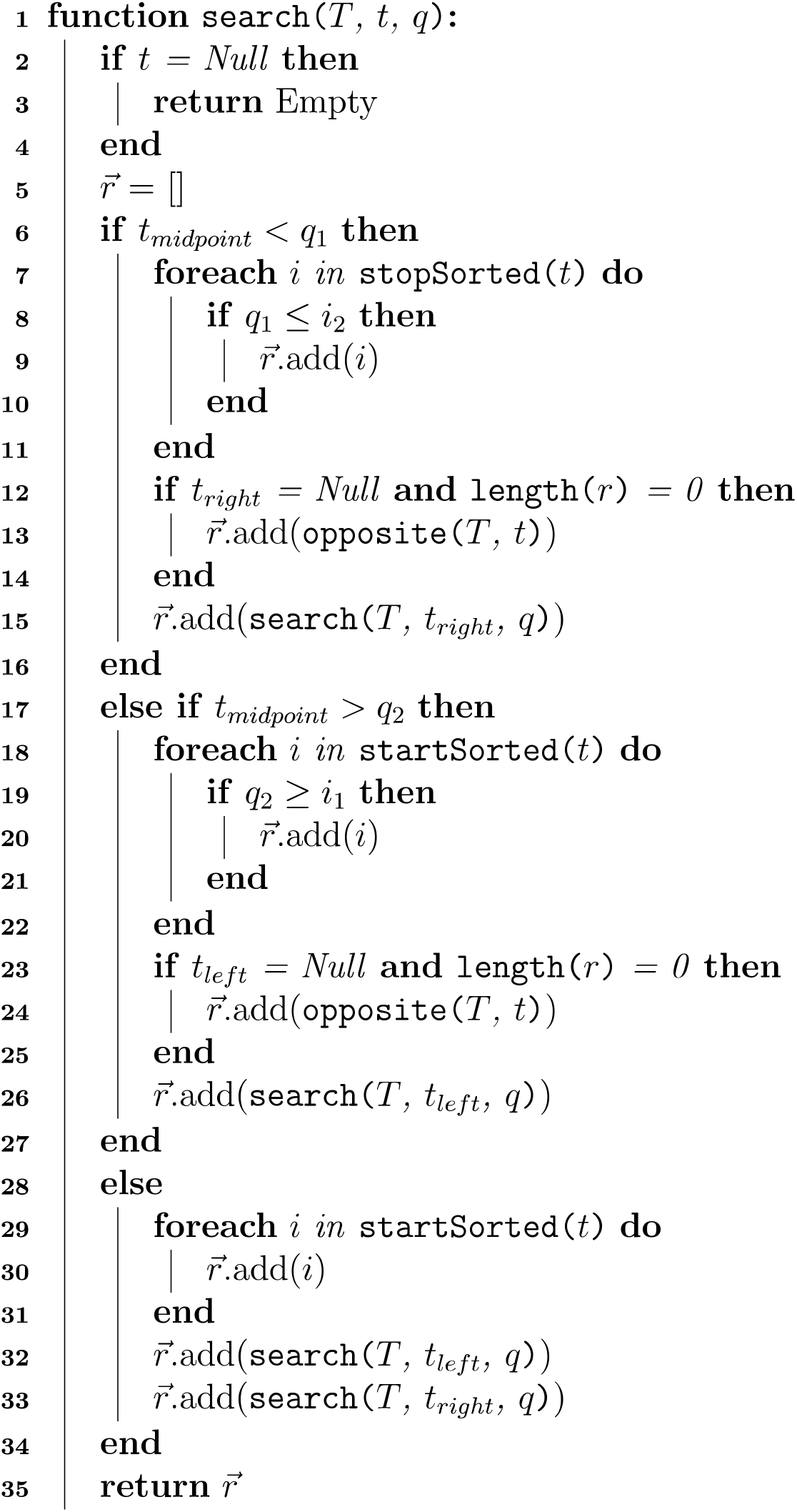
Given the input of a syntenic interval tree (*T*), the current node in the tree (*t*), and a query feature (*q*), find all syntenic intervals in the focal genome that overlap the query feature. 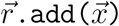 means intervals 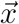 are added to the search result 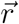. *q*_1_ and *q*_2_ represent the left- and right-hand edges of the query feature. *i* is an interval in the interval tree. *t_midpoint_* is the midpoint of the current node in the tree. *t_1eft_* and *t_right_* and the left- and right-hand subtrees. If no intervals are found, the opposite function returns the nearest blocks on each side of *q* (see **Figure 3**).

### 2.6 Step 5. Map query features and infer search intervals on the target genome

Each query feature is mapped to a search interval that is created with respect to each associated collinear block set (**Figure 2**). To do this, `synder` first classifies each edge of the query feature relative to its relationship to an associated collinear set of blocks (**Figure 4**), where each query feature edge may fall: 1) between two collinear sets of blocks (**unbound**); 2) inside a block (**inblock**); 3) between blocks comprising a collinear block set (**bound**); or 4) beyond all blocks, i.e., near the beginning or end of the scaffold (**extreme**). `synder` sets each boundary of the target genome search interval to the nearest edge of a block in the collinear set if the edge is **inblock** or **bound**; to the nearest syntenic interval beyond the collinear set if the edge is **unbound**; or to the end of the scaffold if the edge is **extreme** (see **Figure 4**).

**Figure 4:**
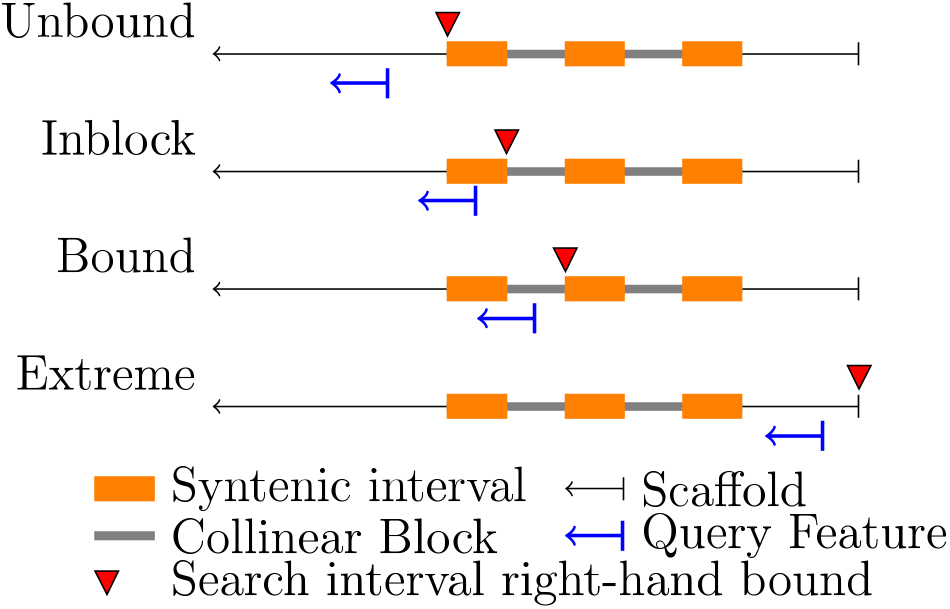
Snapping rules to define the location of the search interval edges on the target genome. 1) The left and right edges of the query feature are used to define the search interval relative to a given collinear block. Only the target genomes and the right edge of the query featured (represented by perpendicular line on query feature) are shown. 2) The right-hand edge of the search interval is then assigned (red triangles) (Rules are the same for the left edge). **Unbound**: the edge does not overlap the collinear block set. **Inblock**: the edge is inside a syntenic interval. **Bound**: the edge is between intervals in a collinear block set. **Extreme**: the edge is beyond any syntenic interval (near end of scaffold).

If a search interval is bound by two blocks, *a* and *b*, which define the two bounding intervals, [*a*_1_, *a*_2_] and [*b*_1_, *b*_2_], then the search interval be the inclusive interval [*a*_2_, *b*_1_]. We use an inclusive interval, rather than the exclusive interval from (*a*_2_ + 1) to (*b*_1_ − 1), to avoid negative length intervals that would occur when *b*_1_ = *a*_1_ + 1 (as would occur if there is a deletion in the target genome).

### 2.7 Step 6. Calculate scores for each collinear set relative to each overlapping query feature

`synder` calculates the *synder score* for each input query feature relative to each associated collinear block set (**Figure 5**). The score reflects the intuitive ideas that: 1) query features are more reliable if a greater proportion of their sequence overlaps blocks in a collinear set; and, 2) query features are more reliable if they are within collinear block sets that are densely packed. In cases where many possible search intervals are identified for a given query feature, the *synder scores* can be used to compare the relative quality of the search intervals. The `synder` score is especially important when *k* (the number of interrupting blocks that is tolerated) is high, since a large *k* allows large gaps between blocks in the collinear block sets. The pseudocode for `synder`’s scoring algorithm is shown in **Algorithm 4**. Note that input scores for syntenic blocks have been transformed to be additive (**Equation 2.2** (Step 1)).

**Figure 5:**
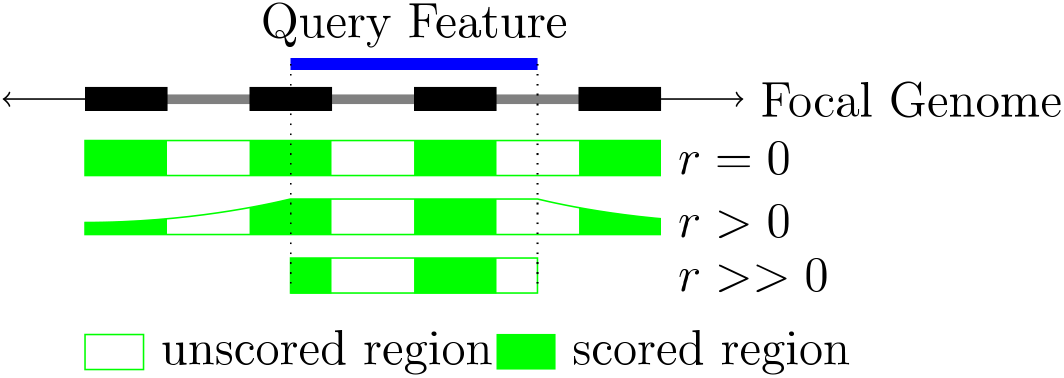
Calculation of `synder` score for a query feature relative to a collinear block set. **Black bars:** the collinear set of syntenic blocks that anchor the focal genome to the target genomes. (Only depicted on focal genome.) The total score of the search interval is the sum of the scores for each block. The score for each block, relative to the query feature (blue bar), is equal to the synteny score for the block times the “weight” of the block (determine by the adjustable parameter, *r*). Three values for *r* are depicted. The weight of each block is the area represented by the **solid green**. The intervals between blocks (**empty green**) do not contribute to the score.

**Algorithm 4:**
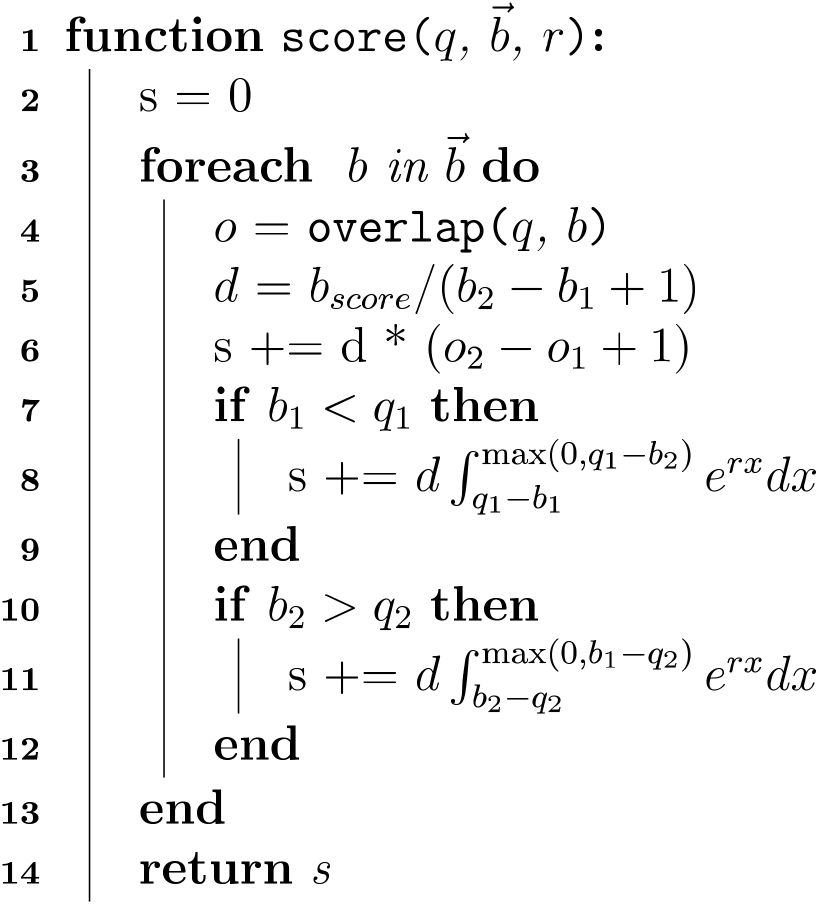
Calculating the *synder score* (*s*) for a query feature and the set of collinear blocks from the target genome to which it is anchored. *q* is the query feature, *b* is a focal genome-side syntenic interval within collinear block set 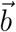, and *o* is the intersection (of zero length or greater) between *q* and *b*. The start and end points (edges) of the query feature *q* are *q*_1_ and *q*_2_ (as for edges of *b* and *o*). *b_score_* is the synteny score associated with syntenic interval *b. r* is an adjustable parameter, the decay rate.

In **Algorithm 4**, each block in the collinear block set can contribute to the total *synder* score (**Figure 5**). The score decay rate is controlled by the adjustable parameter *r*. For the default settings, the weight of the scores of blocks that neither overlap nor partially overlap the query feature decays exponentially with the absolute distance from the nearest query feature bound on the focal side. If the user sets *r* to be a low positive number, the weight at a given position will fall slowly with distance from the query interval (e.g., when *r* = 0.001 the weight will fall by half by 1000 bases from the nearest query feature bound); thus, all blocks in the collinear set will contribute to the score, but they matter less with distance (**Figure 5**, *r* > 0). *r* = 0 would give equal weight to all blocks in the collinear set, in that case, the density of the map will not affect the score, and the score would simply be equal to the sums of the total scores for all the syntenic blocks. A high value, such as *r* = 100, would completely ignore genomic context, basing the query feature score only on the portions of syntenic blocks that overlap the query feature. With this r setting, the `synder` score would be 0 if the query feature does not overlap any syntenic block.

## 3 RESULTS AND DISCUSSION

Mapping genes in a gene family in one species to their orthologs in a related species is a major usage case for `synder`. To demonstrate the use of `synder`, we identify orthologs of the *A. thaliana* NF-YC family genes across several species in Brassicaceae (**Figure 6**).

**Figure 6:**
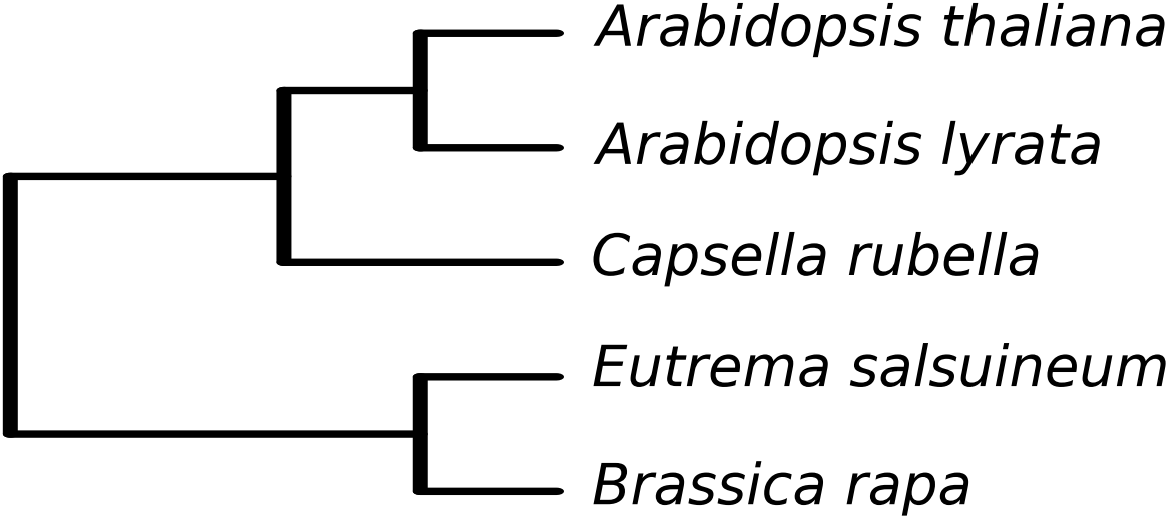
The species tree of the Brassicaceae used in this study. *A. thaliana* is the focal species.

The canonical NF-Y is a heterotrimeric, eukaryotic transcription factor that is comprised of subunits NF-YA, NF-YB and NF-YC [31]. NF-Y has been associated with characteristics as diverse as cell division [32], cancer [33,34], drought tolerance [35], broad spectrum disease resistance [36], and carbon and nitrogen partitioning [37]. In animals and fungi, a single gene encodes each of the three NF-Y subunits. In contrast, the NF-Y subunits in plants are each encoded by gene families of 10-15 genes [34,38,39]. The combinatorial complexity of the potential plant NF-Y complexes that could be formed from the three NF-Y subunits has obfuscated the role of each individual family member, although progress is being made. For example, NF-YA1 of alfalfa controls successful symbiosis between rhizobia and plant, and is required for the persistence of the nodule meristem [40,41].

Compounding the complexity of the action of NF-Y subunits in the cannonical heterotrimer, specific family members of at least two of the subunits, NF-YC and NF-YB, can also form associations with a variety of other nuclear proteins [37,42,43]. One of three NF-YC proteins can interact with one of two NF-YB proteins to enable CONSTANS-promoted photoperiod-induced flowering [44,45]. NF-YC4 of *A. thaliana* interacts with the protein of the orphan gene QQS to modulate carbon and nitrogen partitioning [37]. Several NF-YC family members interact with histone deacetylase 15 (HDA15) in the light to reduce histone acetylation, which in turn decreases hypocotyl elongation [42].

Clear determination of NF-YC orthologs across species would permit the assessment of the relationships among orthology and function in evolution of this gene family. In practice, ortholog identification is often based only on sequence similarity scores. The highest scoring match, however, may not be the true ortholog. `synder` allows more reliable ortholog inference by finding the similarity matches that overlap the inferred syntenic regions. In this way, `synder` may serve as a syntenic filter downstream of the similarity search.

### 3.1 NF-YC orthologs: *Arabidopsis thaliana* compared to *Arabidopsis lyrata*

The specific case of determining the *A. thaliana* NF-YC orthologs in its sister species, *A. lyrata*, illustrates the use of `synder` in resolving orthologs. Since NF-Y is a large family, the paralogous NF-YC family members must be distinguished from the true orthologs.

The genomic relationship between the two species further challenges analysis. While the species diverged only about 8.8 million years ago [46], *A. lyrata* has undergone a whole genome duplication since splitting from the common ancestor it shares with *A. thaliana*. This complicates orthology inference, since each *A. thaliana* gene is expected to have two orthologs in *A. lyrata.* Only one of each duplicate *A. lyrata* ortholog may have preserved a function. Its sister ortholog may have been deleted or become a pseudogene through genome fractionation [28,47]. Alternatively, a sister ortholog may have undergone selection for a completely new molecular function [48,49]. A third possibility is that the molecular function of each sister ortholog was preserved through the neutral process of subfunctionalization [48].

In this analysis, we built a synteny map with Satsuma [20] using default parameters that yielded 229,562 syntenic blocks with a median length of 163 nt (1st quantile = 33 bases, 3rd quantile = 365 bases). This is a very dense map: the *A. thaliana* genome is about 120M in length, thus there are an average of around 1900 syntenic blocks per megabase.

A `synder` search mapped 12 of the 13 query NF-YC family member genes to a search interval in *A. lyrata* that also contained the top BLAST hit (see Hits worksheet in supplementary) (**Figure 7**). Further, `synder` uniquely mapped 11 of the 13 query genes to a single *A. lyrata* gene. NF-YC5 and NF-YC10 are mapped by `synder` to two genes in *A. lyrata*, potentially reflecting that the genome duplication of *A. lyrata* was syntenically conserved. In contrast, a BLAST search yielded a nearly fully connected graph between NF-YC members in the two species. In the case of NF-YC8, `synder` identified orthologs that were located in the same syntenic region of the two genomes; however,the whole genome BLAST did not identify these likely orthologs, but rather other sequences were among the top hits. `synder` and BLAST identified the same gene as having the highest score. A second NF-YC12 ortholog was identified by `synder` that was not selected by BLAST.

**Figure 7:**
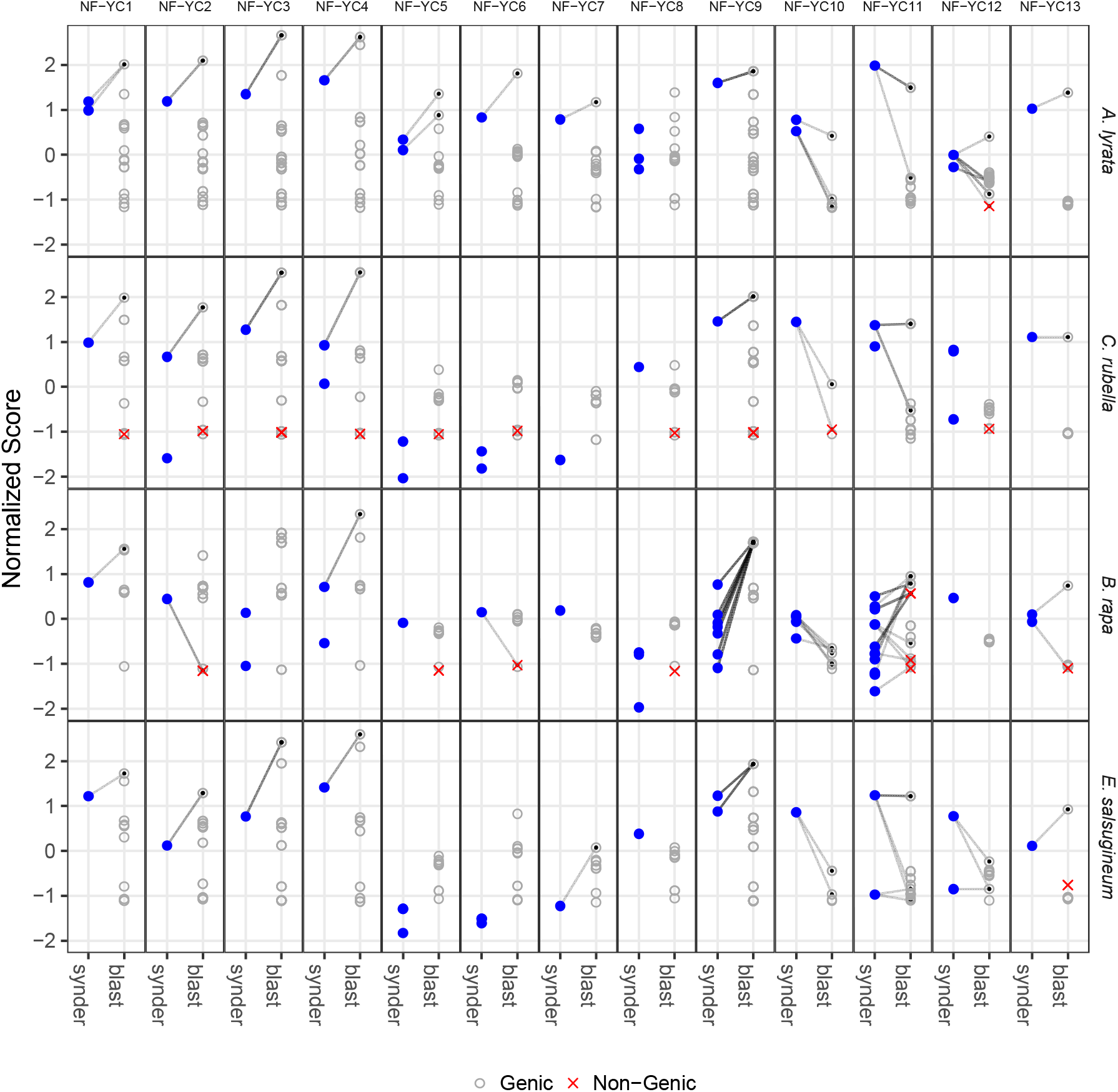
Comparison of orthologs inferred by `synder` and BLAST for the 13 *A. thaliana* NF-YC family members across genomes of four target species from Brassicaceae. Each row represents predicted orthologs in a target species. The x-axis for each box compares `synder` and tBLASTn scores. **Blue dots**, search intervals on the target genome that overlap a gene; **gray circles**, tBLASTn hits (E-value<0.001) on the target genome; **red X’s**, tBLASTn hits on the target genome that do *not* overlap any annotated gene. The normalized Score (y-axis) is the score for `synder` search intervals and tBLASTn hits; `synder` scores were logged, and tBLASTn E-values were transformed with a negated, base10 log. Values were normalized by subtracting the means and dividing by the standard deviation. **Gray lines**, overlap between the tBLASTn hit interval and the `synder` search interval, i.e., tBLASTn finds a hit on the expected strand within the `synder`-inferred search interval.

If each pair of orthologs in *A. lyrata* had undergone only minimal sequence divergence and if synteny was maintained in each case, a `synder` analysis might uniquely identify two *A. lyrata* orthologs for each of the 13 NF-YCs of *A. thaliana.* Indeed, `synder` identified two hits for three of the family member: NF-YC5, NF-YC10 and NF-YC12. The BLAST results cannot reveal whether the second top BLAST hit is an ortholog or not.

### 3.2 NF-YC orthologs across the Brassicaceae family

This approach can be easily extended across the Brassicaceae family. We consider the species in **Figure 6**. In each species, tBLASTn alone links each NF-YC gene to nearly all of the other NF-YC genes; in contrast, `synder` identifies unique mappings to orthologs, many of which differ from the highest BLAST hit. As syntenic distance increases, the orthologs become more difficult to identify through syntenic methods (**Figure 7**).

However, for those genes located in syntenically-conserved regions, `synder` would more reliably identify the ortholog. For example, in *B. rapa*, `synder` identifies an NF-YC ortholog within the syntenic search space for NF-YC3, NF-YC5, NF-YC7, and NF-YC8 that does not correspond to the top BLAST hit (**Figure 7**). `synder` identified a syntenic ortholog of NF-YC6 that is only a weak BLAST hit. For NF-YC2, the ortholog identified by `synder` is not annotated at all in *B. rapa.* It may or may not be an expressed gene, but it is most likely an ortholog. Thus, `synder` can be used to augment whole-genome similarity inferences with syntenic context information.

## 4 CONCLUSION

`synder` provides a flexible, reproducible method to track specific genetic events. It also provides a pathway to evaluate broad biological concepts, including the evolution and diversification of gene families; the predominant mechanisms of diversification across lineages of eukaryotes and prokaryotes; the effects of genome duplication; and the relationship of different features to genomic instability. These types of analyses can ultimately reveal those genetic events that might be associated with particular evolutionary consequences, such as rapid evolution, horizontal transfer, *de novo* emergence of genes, transposition, or duplication.

## 5 IMPLEMENTATION

`synder` is a C++ program wrapped in an R package via Rcpp [50]. It is designed to be compatible with Bioconductor, an R-based bioinformatics ecosystem [51].

## 6 AVAILABILITY

As an R package, `synder` should work on any system. It is distributed under a GPL-3 open source license and the source code is available at https://github.com/arendsee/synder. All code required to run the case study is available at https://github.com/arendsee/synder-case-study and the required input data is on DataHub at https://datahub.io/arendsee/synder-nfyc.

## 7 ACKNOWLEDGEMENTS

We thank Steven Cannon and Jennifer Chang for discussion and proof-reading.

## 8 AUTHORS’ CONTRIBUTIONS

ZA conceived of the project and ZA and AW implemented the software. MH reviewed the software and offered helpful feedback. US and LJ tested the software and helped with the case studies. ZA and ESW wrote the initial drafts and final version. All authors participated in the writing of the manuscript, and read and approved the final manuscript.

